# An ultra high-throughput, massively multiplexable, single-cell RNA-seq platform in yeasts

**DOI:** 10.1101/2022.09.12.507686

**Authors:** Leandra Brettner, Rachel Eder, Kara Schmidlin, Kerry Geiler-Samerotte

## Abstract

Yeasts are naturally diverse, genetically tractable, and easy to grow in a myriad of experimental conditions such that researchers have the ability to investigate any number of genotypes, strains, environments, or the interaction thereof. However, studies of variation in the yeast transcriptome have been limited by the processing capabilities of available RNA sequencing techniques. Here we optimize a powerful, high-throughput single-cell RNA sequencing (scRNAseq) platform for yeasts. This platform utilizes a combinatorial barcoding strategy to enable massively parallel RNA sequencing of hundreds of yeast genotypes or growth conditions at once. This method can be applied to most species or strains of yeast for a fraction of the cost of traditional scRNAseq approaches. Thus, our technology permits researchers to leverage “the awesome power of yeast” by allowing us to survey the transcriptome of hundreds of strains and environments in a short period of time, and with no specialized equipment. The key to this method is that sequential barcodes are probabilistically appended to cDNA copies of RNA while the molecules remain trapped inside of each cell. Thus, the transcriptome of each cell is labeled with a unique combination of barcodes. Since we use the cell membrane as a container for this reaction, many cells can be processed together without the need to physically isolate them from one another in separate wells or droplets. Further, the first barcode in the sequence can be chosen intentionally to identify samples from different environments or genetic backgrounds, enabling multiplexing of hundreds of unique samples in a single experiment. In addition to greater multiplexing capabilities, our method also facilitates a deeper investigation of biological heterogeneity given its single-cell nature. For example, in the data presented here we report transcriptionally distinct cell states related to cell cycle, growth rate, metabolic strategies, stress responses, etc. all within clonal yeast populations grown in the same environment. Hence, our technology has two obvious and impactful applications for yeast research: the first is the general study of transcriptional phenotypes across many strains and environments, and the second is investigating cell-to-cell heterogeneity across the entire transcriptome.

## INTRODUCTION

Yeasts are probably one of the most well-researched lifeforms, and serve as workhorse model organisms for exploring genetics, evolutionary and systems biology. They are arguably the most genetically tractable eukaryote with the easy construction of comprehensive gene deletion collections (Giaever et al., 2002), and more recently, libraries of thousands of strains in a single genetic editing experiment (Levy et al., 2015; Sharon et al., 2018). There is also a wealth of well-annotated, natural genetic diversity available for study (Liti et al., 2009; Peter et al., 2018). However, measuring the phenotypic effects of that many mutations is largely unfeasible. Further, to fully survey all of those genotypes, each strain should ideally be assessed in multiple conditions to grasp the effects of environmental context. This quickly multiplies to sample numbers that are hard to manage.

Single-cell RNA sequencing (scRNAseq), or the profiling of RNA expression in individual cells, has become a powerful tool that enables the high-throughput interrogation of transcriptional phenotypes across multiple strains and multiple conditions simultaneously (Dixit et al., 2016; Jackson et al., 2020; Rodriguez-Fraticelli et al., 2020). While several recent papers have been published applying scRNAseq to yeast, these papers utilize scRNAseq methods involving physical isolation of cells and accumulate expense both in the specialized equipment and the number of cells processed (Kolodziejczyk et al., 2015; Liu and Trapnell, 2016). Recently developed combinatorial barcoding methods (SPLiT-seq (Rosenberg et al., 2018), sciSeq (Cao et al., 2017)) solve these issues by being easily scalable and only requiring basic benchtop tools. However, to date, these methods have been applied to a variety of organisms (Cao et al., 2017; Kuchina et al., 2021; Rosenberg et al., 2018), but, to our knowledge, not to fungi. Here we present a yeast optimized version of Split Pool Ligation-based Transcriptome sequencing, or SPLiT-seq (Kuchina et al., 2021; Rosenberg et al., 2018).

This method is more desirable than other available methods because it allows for the sequencing of the transcriptomes of far more cells, giving us more power to profile many strains and conditions as well as phenotypic heterogeneity. For example, our method can process ~400,000 cells for less than $2,000 (Rosenberg et al., 2018), while droplet-based methods would cost >$40,000 to do the same based on price listings on the University of Kansas Medical Center, Boston University Medical Center Sequencing Core, and Cornell Institute of Biotechnology websites. Note that this calculation does not include the large cost of the of the instrument but assumes that researchers are submitting their samples to sequencing cores or companies. And our method can include as many samples as there are barcodes (eg. 96 or 384) per experiment depending on the type of multi-well plates used, while droplet-based methods can handle up to only 12 samples per run (10X Genomics 3’ CellPlex Kit).

## TECHNIQUE OVERVIEW

In opposition to isolation-based scRNAseq methods that utilize microfluidic droplets to contain the RNA from a single cell (Dohn et al., 2021; Macosko et al., 2015; McNulty et al., 2021; Zheng et al., 2017), the SPLiT-seq protocol uses the cell itself as a container for its own RNA, and all enzymatic reactions are performed *in situ* (Figure 1, (Kuchina et al., 2021; Rosenberg et al., 2018)). Fixed and permeabilized cells are loaded into a multi-well plate where they undergo *in situ* reverse transcription with well-specific barcoded random hexamer and poly-T primers. They are then pooled and split into another 96-well plate where each well contains a short, unique barcode sequence that anneals to the first barcode via a staple strand. A ligation reaction covalently bonds these two pieces of DNA at the single stranded nick created. The cells are subsequently pooled and split into a new plate where the process is repeated, adding a third barcode. Cells that received the same first barcode are unlikely to receive the same second and third barcodes. This process is completed n times depending on the population size, as unique barcode combination possibilities scale exponentially with each additional round (e.g. 96 barcodes, n split-pools = 96_n_ possible barcode combinations). Each cell is thus uniquely labeled by probabilistically biasing the outcome such that it takes its own path through the barcode plates. Finally, the cells are lysed, and the extracted cDNA is prepped for sequencing (Figure 1). After sequencing, combinatorial barcodes are used to computationally resolve through which wells a cell has traveled in order to get single-cell data (Kuchina et al., 2021; Rosenberg et al., 2018).

**Figure 1:**
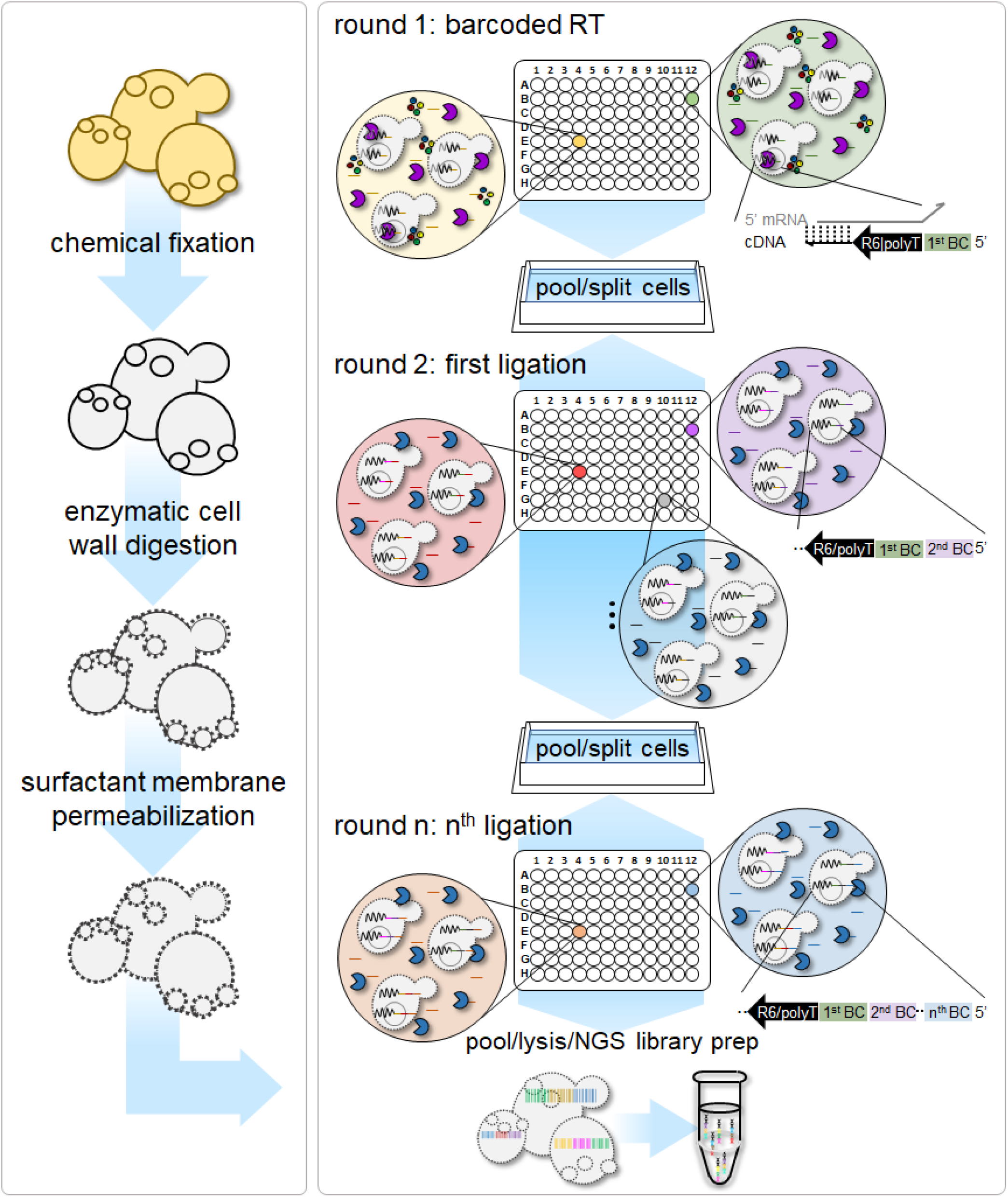
Yeast optimized SPLiT-seq. **Left panel)** Yeast specific protocols were developed for the fixation, cell wall enzymatic digestion, and surfactant membrane permeabilization steps. These are the steps that are the most important to adapt to the working organism, as each creature is going to have different structural morphologies that will need customization, particularly for the permeabilization steps. For example, yeasts have cell walls whereas mammalian cells do not. **Right panel)** The split-pool reverse transcription and ligation steps are carried out in multi-well plates similar to the original protocols (Kuchina et al., 2021; Rosenberg et al., 2018). The resulting cNDA libraries are prepared and submitted for sequencing.

The intrinsic power of SPLiT-seq comes from its easy and inexpensive capabilities to scale up the number of cells processed. The cost of single-cell sequencing is roughly correlated to the number of containers used. In physical isolation methods (e.g., droplet methods), every cell needs its own container in order to label its transcriptome with a unique barcode. Thus, adding more cells means adding a proportional number of new containers. However, the SPLiT-seq method does not scale in the same linear way (Figure 2a). The number of cells that can be processed is a function of the number of wells in each barcoding step and the number of ligation reactions performed (Figure 2a). For example, one round of barcoded reverse transcription with 96 barcodes and two subsequent ligation steps, also with 96 barcodes, produces 96^3 (884,736) possible unique combinations. Adding one more ligation reaction yields almost 85 million unique combinations with only the trivial cost of adding another plate of 96 barcode oligos and reagents (Figure 2a).

**Figure 2:**
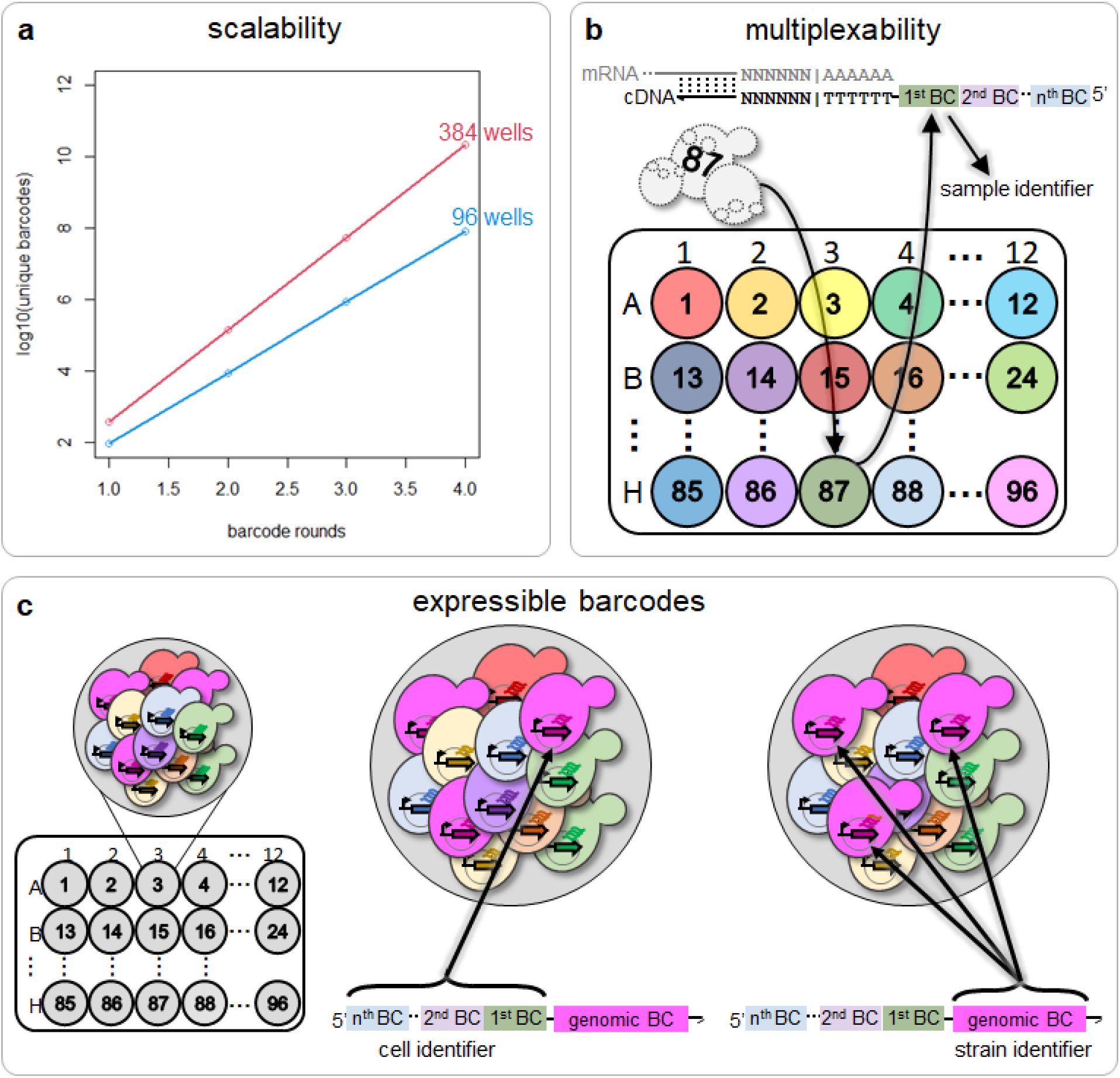
The incredible high-throughput power of the SPLiT-seq method. **a)** The number of available unique barcode combinations scales exponentially with addition of relatively few new barcodes (increments of 96 or 384). A similar scaling analysis can be found in the original SPLiT-seq publication (Rosenberg et al., 2018) **b)** The number of samples that can be processed in a single run is equal to the number of barcoded primers in the reverse transcription step, which provides the first barcode in the combinatorial sequence. For example, 96 or 384 samples could be processed with standard multi-well plates. **c)** Expressible barcodes, or expressed engineered sequences that identify genotype, can be combined with the SPLiT-seq method to provide incredible genotype and environment combinatorial power.

Another advantage of SPLIT-seq is the ability to multiplex many samples in the same run. Since each sample can be loaded intentionally into a specified well of a 96- or 384-well plate during the reverse transcription step, the first barcode in the combinatorial sequence can be used as a conditional signifier. This allows for the processing of as many unique samples as there are wells in the first step (Figure 2b). So, for example, if a 96 well scheme is used, 96 samples representing 96 genotypes or 96 environments can be processed in that run. Using expressible barcodes (Figure 3c, (Jackson et al., 2020)) to label unique genotypes can increase the number of samples even further, enabling massive GxE screens. Assuming each strain’s transcriptome requires 500 cells per experiment to achieve adequate coverage, a single standard SPLiT-seq run (3 rounds of 96 barcodes) could process almost 900,000 cells. This is enough to cover as many as 1800 genotypes if expressible barcodes are used, or 1800 different combinations of genotype and environment. As each SPLiT-seq run takes less than a week to perform, hundreds of conditions and thousands of strains could easily be sampled in only a few months. Additionally, as cells are fixed right after sampling, early samples can be stored in the freezer until all experiments are finished and ready for scRNAseq preparations.

**Figure 3:**
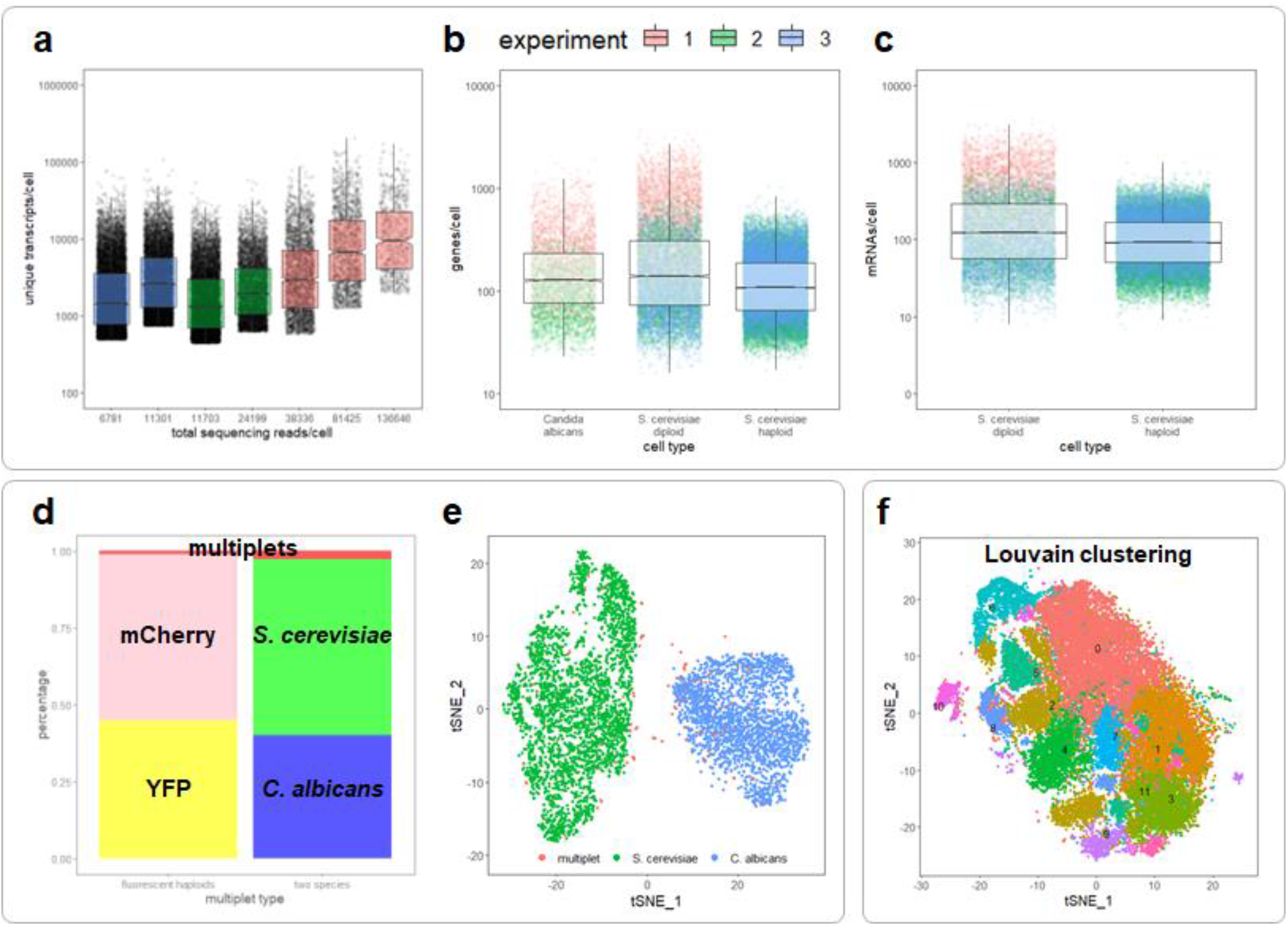
Yeast optimized SPLiT-seq produces quality scRNAseq data. **a)** Total unique transcripts detected per cell (including rRNAs) for multiple sublibraries across 3 experiments. Each point represents a single cell and boxes represent 1st - 3rd quartiles. Median transcripts per cell range from1500 to 9500 per cell. When fewer cells are profiled (rightmost sublibraries), the sequencing depth per cell and the number of unique transcripts per cell increase. We use unique molecular identifiers to distinguish unique transcripts from duplicates created during PCR. **b)** Total genes per cell detected in *C. albicans*, and diploid and haploid *S. cerevisiae*. Box represents 1st - 3rd quartiles. **c)** Total mRNAs per cell detected in diploid and haploid *S. cerevisiae*. Box represents 1st - 3rd quartiles. **d)** Percentages of barcodes that classify as YFP or mCherry expressing cells, *S. cerevisiae* or *C. albicans*, or multiplets (barcodes that have any reported expression of the opposing fluorescent reporter, or greater than 15% of the opposing species’ genome) **e)** tSNE visual clustering of *S. cerevisiae* and *C. albicans* data. Each point represents a single barcode or cell and its position relative to other points correlates with the similarity of their transcriptomes. The points are colored by their mapped or “true” identity. We see a distinct separation of the two species. **f)** tSNE visual clustering of combined haploid and diploid S. cerevisiae cells. The number-labeled, colored clusters represent clusters detected using a standard Louvain algorithm in Seurat (Hao et al., 2021).

In sum, SPLIT-seq is an incredibly high-throughput method for single-cell RNA sequencing that lends itself to the yeast system because there are so many engineered and natural yeast genotypes to explore. Here we optimize the method for the unique physiology of yeasts and present data demonstrating its effectiveness.

## RESULTS

### Yeast optimized SPLiT-seq detects thousands of unique RNAs per cell for a variety of yeast ploidies and species

After modifying the protocol to work with yeasts, SPLIT-seq detects a median of ~1000 to 10,000 unique RNAs per cell depending on sequencing depth. Higher sequencing coverage results in a deeper sampling of the transcriptome as can be seen in Figure 3a which shows median unique RNAs per cell for technical replicates of three experiments of different sequencing depths. It is effective in both haploid and diploid cells of *S. cerevisiae*, and also works with *C. albicans*. We recover, on average, between 100 and 600 genes per cell (Figure 3b). Of the detected RNAs, we see an average of ~120 – 700 per cell depending on the experiment and sequencing depth. Across all cells, we see the range of detected mRNAs being between ~10 to 3600 per cell (Figure 3c). Given that there are approximately 30,000 mRNA molecules per cell for *Saccharomyces cereivisiae* (Miura et al., 2008), and assuming haploid and diploid *S. cerevisiae* and *C. albicans* have transcriptome sizes in the same order of magnitude, this range represents 0.2-12% of the active transcriptome in a yeast cell.

### Yeast optimized SPLiT-seq is truly single-celled: it experiences a negligible percentage of barcode collisions

A common concern in single-cell RNA sequencing experiments is barcode collisions, or one barcode mapping to multiple cells. In physical isolation methods, this occurs when more than one cell is loaded into the same container. In the SPLiT-seq method, multiple cells can acquire the same barcode by physically aggregating or through cells traveling the same path through the barcode plates by chance. We performed two experiments to get an empirical estimate of the percentage of barcodes that map to multiple cells, or multiplets. We first processed two disparate species, diploid *S. cerevisiae* and *C. albicans*, mixed together in the same SPLiT-seq run. We could then calculate the percentage of barcode collisions by calculating the number of barcodes that have a large percentage (>15%) of uniquely mapping transcripts that align to both of the species’ genomes. We removed rRNA reads before this calculation as there are considerable amounts of homology between these species in those genes and keeping them in falsely increased the number of calculated barcode collisions.

Since many engineered yeast strains are haploid *S. cerevisiae*, we also wanted to ensure that these cells did not have any odd properties that caused them to stick together. We looked at stickiness between haploid *S. cerevisiae* by using two haploid strains each expressing an engineered fluorescent protein. We looked for collisions by counting the number of barcodes that report expression of both fluorescent markers. Figure 3d shows that in both experiments, the percentage of detected barcode collisions is well under five percent. We also see good species resolution in tSNE based clustering analyses (Figure 3e). This suggests that our method is truly single celled.

### Yeast optimized SPLiT-seq identifies heterogeneous transcriptional states in clonal populations grown in similar conditions

A good test of any single-celled method is whether it can resolve transcriptionally distinct states, for example, those related to the cell cycle. All of the cells in the above experiments were grown in rich media and sampled in early mid log phase. Even in this carbon rich environment, we see considerable diversity in gene expression (Figure 3f). We performed the Louvain clustering algorithm to group cells based on the gene expression data we collected and used tSNE to visualize the cell clusters. For each of these clusters, we performed a differential gene expression analysis to investigate what biological processes were distinguishing cells based on their gene ontology as assessed using Metascape and the Saccharomyces Genome Database (Cherry et al., 2012; Zhou et al., 2019). For example, cluster 2 appears to be cells which have very recently divided, as the top differentially expressed genes are related to septum digestion after cytokinesis. It might be more specifically newly divided daughter cells as we also observe several daughter-specific upregulated genes such as DSE1, DSE2, and DSE4. Cells in cluster 4 appear to be in late G1 or early S phase as we see upregulated expression of the G1 cyclin CLN2, genes involved in the formation of the bud neck such as HSL1 and GIN4, and genes related to DNA replication such as POL1 and RNR1. A full list of statistically differentially expressed genes after multiple comparison correction is available in Supplementary Table 1. These data suggest that our method is sampling the transcriptomes of individual cells with enough depth to cluster cells based on their transcriptionally distinct states.

## DISCUSSION

The number of mRNAs we recover using our yeast-optimized SPLiT-seq method (120-700 on average per cell) and the number of genes (100-600 on average per cell) is in the same ballpark but slightly less than reported recovery for yeast experiments performed with the 10X genomics microfluidics based platform (~2000 mRNAs per cell and ~700 genes per cell (Jackson et al., 2020)). Nonetheless, there are several reasons our method is likely to be useful to the yeast community. First, an average of hundreds of unique reads per cell is enough for many applications. For example, we recover enough data to delineate biologically relevant cell states (Figure 3f). Additionally, published studies detect less than 5% of each cell’s transcriptome, but by clustering similar cells, are able to comprehensively map different transcriptional states (Kuchina et al., 2021). Second, we observe up to thousands of mRNAs in some cells, suggesting further optimization may yet yield further improvements. And finally, we believe the highly scalable and massively multiplexable nature of the technique make it an important addition to the yeast community. If any researchers would like to try the yeast optimized SPLiT-seq method, we are happy to correspond about experimental design and share detailed protocols.

## METHODS

### Experimental Methods

#### Yeast cell culture

Diploid s288c, genetically naive haploid BY4741, *MATa ura3A0 his3*Δ *1 met17*Δ *0 P ACT1-GAL3::SpHIS5 gall1*Δ *gal10*Δ *::LEU2 leu2*Δ *0::P PGK1 -mCherry-KanMX6 ybr209w*Δ *::B103- HphMX6*, and *MATa ura3A0 his3Δ 1 met17*Δ *0 P ACT1-GAL3::SpHIS5 gall1*Δ *gal10*Δ *::LEU2 leu2*Δ *0::P GAL1* -*YFP-KanMX6 ybr209w*Δ *::BC3-HphMX6*, and ATCC 3147 *C. albicans* strains were used. Cells were streaked on YP plus 2% dextrose agar plates from frozen -80C glycerol stocks and grown at 30C for 48 hours. In experiments 1 and 2, single colonies of s288c and ATCC 3147 were picked into YP plus 2% dextrose liquid media. The YFP and mCherry engineered cells were picked into Synthetic Complete minus glucose (Sunrise) plus 2.5% sucrose, 1.25% raffinose, and 0.625% galactose media to induce YFP expression in experiment 2. In experiment 3, diploid s288c cells and haploid BY4741 cells were picked into Synthetic Complete (Sunrise) plus 2% glucose media. All cultures were grown with shaking at 30C for 24 hours. Cells were then transferred into fresh media at a 1:250 dilution and grown until cultures reached approximately 2-4*10^7 cells/mL, or early mid log phase.

#### Fixation and permeabilization

At the time of sampling, 3 mL of yeast cultures were immediately spun down in a room temperature centrifuge at 3000G for 5 minutes. The media supernatant was removed and the cell pellet was resuspended in 2 mL of cold 4% formaldehyde in molecular grade PBS. The cells were then fixed cold in a 4C refrigerator for approximately 18 hours. After fixation, the samples were spun down in a 4C centrifuge at 3000G for 5 minutes. The formaldehyde supernatant was removed and the pellet was resuspended in 1 mL cold molecular grade 100 mM Tris HCl pH 7 plus 0.1 U/uL SUPERase-In RNase inhibitor (ThermoFisher) to quench the formaldehyde. Cells were then centrifuged, resuspended in 250 uL of a zymolyase enzymatic solution of 0.1M Na_2_EDTA, 1M sorbitol, and 0.01 U/uL zymolyase (Zymo Research) (pH ~7.5), and incubated at 37C for 15 minutes. At the end of the incubation, 1 mL of cold PBS plus 0.1 U/uL SUPERase and 0.1 U/uL Enzymatics RNase Inhibitor (Enzymatics) was immediately added and the cells were centrifuged at 4C, 3000G for 5 minutes. The cells were then resuspended in 250 uL of cold 0.04% Tween-20 in PBS plus RNase inhibitors and incubated on ice for 3 minutes. Again, 1 mL of cold PBS plus RNase inhibitors was added at the end of the 3 minutes and the cells were centrifuged. They were then resuspended in 500 uL of cold PBS plus RNase inhibitors and counted using a Beckman Coulter Cell Counter. The cells were then diluted to 1 million cells/mL into fresh cold PBS plus RNase inhibitors.

#### *In situ* reverse transcription, *in situ* ligation, and lysis

were performed as described in previous work (Kuchina et al., 2021).

#### cDNA purification, template switch reaction, and bead PCR

were performed as described in previous work (Rosenberg et al., 2018).

#### WGS fragmentation library preparation

was performed as described in previous work (Kuchina et al., 2021).

#### Illumina sequencing

Sequencing libraries from each of the 3 experiments were submitted to Psomagen. Experiment 1 was paired end sequenced on a full lane of the HiSeq X platform with a 6 basepair index on read 2. Experiments 2 and 3 were also sequenced on full lanes of the HiSeq X platform, but with dual 8 bp indices.

#### Oligonucleotides

All oligonucleotide sequences used are available in Supplementary Table 2.

### Computational Methods

#### Transcriptome alignment

Sequencing reads were processed by barcode and aligned to the R64 s288c genome or a combined R64 s288c and sc5314 V4 genome from NCBI using STARsolo (Kaminow et al., 2021). This new version of STAR parses out cell barcodes and allows for the multi-mapping of reads, meaning it keeps reads that map to multiple places in the genome. This is important for yeast as *S. cerevisiae* recently underwent a full genome duplication and there is significant homology between many paralogs (Wolfe, 2015). We used the most basic uniform multi-mapping algorithm. This produced a highly sparse gene-by-cell matrix with cell columns for every possible barcode combination, regardless of gene detection.

#### Data quality thresholding

To remove empty barcodes and low gene detection barcodes, we applied an “knee” detection filtering to the STARsolo generated gene-by-cell matrix (Kaminow et al., 2021). Briefly, the barcodes were negatively ordered by log(total read counts). We remove any barcodes after the curve begins to drastically decrease, or any barcodes past the bend or the “knee”. Depending on the library quality, this keeps between 60-80% of the non-zero barcodes.

#### Single-cell data analysis

All data analysis was performed with the R based package, Seurat (Hao et al., 2021). We performed normalization, scaling, nearest neighbor calculations, clustering, and differential expression analyses similar to the tutorial provided by the Satija Lab (“Getting Started with Seurat,” n.d.). Unique transcripts/cell and genes/cell were calculated with all genes left in the data. However, mRNAs/cell, Louvain clustering, and tSNE visualization were done on data with the ribosomal RNA removed. rRNA genes were identified using the Saccharomyces Genome Database (SGD (Cherry et al., 2012)).

#### Gene ontology analysis

Simple gene ontology analyses were done by manually entering genes into both Metascape and SGD (Cherry et al., 2012; Zhou et al., 2019).

## ACKNOWLEDGEMENTS

We would like to thank Yuichi Eguchi and Pak Poon for their helpful insights into yeast morphology, physiology, and culturing methods. Funding for this work was provided by a National Institutes of Health grant R35GM133674 (to KGS), an Alfred P Sloan Research Fellowship in Computational and Molecular Evolutionary Biology grant FG-2021-15705 (to KGS), and a National Science Foundation Biological Integration Institution grant 2119963 (to KGS). We would also like to thank the members of the Geiler-Samerotte Lab for their support and feedback.

